# Exploring C1 substrate cofeeding in *Eubacterium limosum* with AneVO, a low-cost anaerobic parallel bioreactor platform

**DOI:** 10.1101/2025.05.02.651948

**Authors:** Kathryn O. Hoyt, Guanyu Zhou, William Gasparrini, Patrick J. Sliter, Daniel J. Hart, Ahmad S. Khalil, Benjamin M. Woolston

## Abstract

Acetogenic bacteria have emerged as attractive biocatalysts for renewable biochemical production, using the highly efficient Wood-Ljungdahl pathway to convert a range of sustainable single-carbon (C1) feedstocks. The major challenge is their energy-constrained anaerobic lifestyle, which results in slow growth and limits the product spectrum. To overcome this limitation, here we investigate substrate co-metabolism in the acetogen *Eubacterium limosum*, cofeeding either carbon monoxide (CO) or glucose alongside the primary C1 substrate (methanol or formate). To increase experimental throughput, we developed AneVO, a parallel mini bioreactor system based on eVOLVER that enables benchtop anaerobic batch and fed-batch cultivation, along with continuous delivery of anaerobic gas blends. With all substrate pairs tested, *E. limosum* grew faster and reached 52-254% higher cell densities with cofeeding, while maintaining or improving the volumetric uptake rate of the main C1 substrate. Product formation also improved, with an increase in volumetric acetate productivity from glucose cofeeding of 2.2-fold with formate and 2.4-fold with methanol, and 3-fold from CO cofeeding with methanol. Together these results validate AneVO as a low-cost platform for convenient benchtop cultivation of strict anaerobic microbes in multiple growth modes, and present a strategy for enhancing C1 bioconversion rates in *E. limosum*, an emerging model acetogen.

## Introduction

Transitioning fuel and chemical production to microbial systems that use sustainable carbon feedstocks is an essential component of decarbonizing the economy and reducing the harmful environmental impact of the current fossil fuel-based paradigm (Lee *et al*., 2019; Aggarwal *et al*., 2023; Jones *et al*., 2024). Single-carbon (C1) feedstocks, including CO, formic acid, methanol, and methane, have emerged as attractive renewable substrates for bioprocessing (Baumschabl *et al*., 2024). Syngas-like mixtures containing CO, CO_2_ and H_2_ can be produced from biomass or municipal solids waste via gasification, and are also available directly in industrial waste streams such as steel mill off-gas (Daniell *et al*., 2012; Liew *et al*., 2016). Concentrated CO_2_ streams can also be converted into the liquid intermediates formic acid and methanol by electrochemical methods or with renewable H_2_, enabling easier storage and transport (Liew *et al*., 2016; Jouny *et al*., 2018; Cotton *et al*., 2020; Bachleitner *et al*., 2023). These supply chains are completely decoupled from crop production, avoiding food vs. fuel and land use concerns raised over traditional carbohydrate feedstocks (Dürre and Eikmanns, 2015).

Acetogens are especially promising microbial biocatalysts for C1 bioconversion, as they natively metabolize CO_2_, CO, formate and methanol with very high energetic efficiency and carbon yield (Claassens *et al*., 2019). These anaerobic bacteria grow using the Wood-Ljungdahl Pathway (WLP or reductive Acetyl-CoA pathway) and convert as much as 90% of the consumed carbon into acetate and ethanol (Fast and Papoutsakis, 2012). Acetogenic syngas fermentation for production of ethanol from steel mill off-gas (a mixture of CO_2_, CO, and H_2_) has been commercialized, with promising life cycle assessment metrics (Liew *et al*., 2022).

Though less studied than gas fermentation, formate and methanol are also used by certain acetogens, again with exceptionally high energy efficiency (Claassens *et al*., 2019; Cotton *et al*., 2020). Acetate production from methanol via the WLP results in simultaneous consumption of CO_2_, leading to greater than 100% carbon yield (Kremp *et al*., 2018). Methanol provides more ATP per acetate than syngas fermentation (Dietrich *et al*., 2021), as well as additional reducing power which enables cells to produce higher value products such as butyrate (Litty and Müller, 2021; Wood *et al*., 2021). When grown on formate, acetate is typically the exclusive product (Litty and Müller, 2021; Moon *et al*., 2021; Wood *et al*., 2021). In addition to higher yield, acetogens show competitive specific substrate uptake rates when compared to their aerobic counterparts (Claassens *et al*., 2019). At industrial scale, anaerobic processes also benefit from reduced cooling and aeration costs, which are particularly high in aerobic C1 processes (Frazão and Walther, 2020). Finally, the systems and synthetic biology toolboxes for acetogens have expanded dramatically in the last decade, facilitating metabolic engineering campaigns for non-native products (Liew *et al*., 2022; Sanford and Woolston, 2022b; Poehlein *et al*., 2024). Despite the advantages of acetogens, there is one major downside – the limited availability of cellular energy during anaerobic C1 fermentation (Molitor *et al*., 2017). For each of the C1 substrates discussed, the WLP produces less than one mole of ATP per mole of acetate (Schuchmann and Müller, 2014; Cotton *et al*., 2020). This results in long doubling times and low cell densities, which together limit volumetric productivity, increasing reactor size and capital cost. It also limits the yield of non-native products, since the cells must continue to produce acetate as a byproduct in order to balance their ATP needs (Fast and Papoutsakis, 2012). Indeed, the list of target products that have been made in acetogens with competitive metrics is short (Liew *et al*., 2022).

One means to increase energy availability and improve growth on C1 substrates is cofeeding, or mixotrophy (Fast *et al*., 2015; Liu *et al*., 2020; Lee *et al*., 2022). In this paradigm, a second carbon source is added to the fermentation in a manner that enables simultaneous consumption of both substrates, resulting in faster growth. For example, in the thermophilic acetogen *Moorella thermoacetica,* controlled glucose feeding during growth on CO_2_/H_2_ led to a 4- fold increase in the specific CO_2_ uptake rate compared to fully autotrophic growth, reaching an impressive 2.3 g CO_2_ gCDW^-1^ h^-1^ (Park *et al*., 2019). In the methylotrophic acetogen *Eubacterium limosum,* cofeeding a mixture of glucose and methanol resulted in higher biomass yields and specific growth rates than on either substrate alone (Loubiere *et al*., 1992). Substrate co-feeding can also positively impact product yield. Mixotrophic growth on syngas and fructose led to an exceptionally high acetone yield in an engineered strain of *C. ljungdahlii,* reaching 138% higher than the theoretical maximum on fructose alone (Jones *et al*., 2016).

Beyond mixtures of sugars and a C1 substrate, cofeeding in acetogens has also been explored with combinations of C1 substrates. A recent study with a butyrate-producing *Acetobacterium woodii* co-metabolizing methanol and CO reported improved butyrate titers compared to methanol alone, likely due to the availability of more reducing power from CO (Chowdhury *et al*., 2022). CO and formate cofeeding also led to growth and productivity improvements in *A. woodii* and *Clostridium ragsdalei* (Neuendorf *et al*., 2021; Schwarz *et al*., 2024). In some cases, such combinations lead to formation of new metabolites. Co-metabolism of formate and methanol in *E. limosum* results in production of butanol, which is not detected during growth on either substrate alone (Wood *et al*., 2021, 2023). The emergence of atypical metabolites was also observed in the closely related *Butyribacterium methylotrophicum*, where methanol/CO cofeeding lead to production of butanol, ethanol, and lactate (Humphreys *et al*., 2022).

These previous studies clearly establish substrate co-metabolism as a general phenomenon in acetogens for improving growth and/or production on C1 substrates. The mechanisms underlying the growth benefits, however, are not always clear. In *E. limosum,* for example, the reported benefit of glucose cofeeding during growth on methanol could be attributed to the extra ATP produced from glycolysis compared to the WLP (Loubiere *et al*., 1992). Another possibility is that the second substrate enables the cell to produce critical biosynthetic metabolites through pathways that are less thermodynamically or kinetically challenging, or more optimally balance the use of different energy cofactors (Le Bloas *et al*., 1993). Delineating between these possibilities is critical to inform metabolic engineering strategies to optimally exploit cofeeding in acetogens engineered to produce high-value non-native products (Liu *et al*., 2020). Further, an understanding of the mechanism by which cofeeding improves growth on C1 substrates could identify specific metabolic bottlenecks that could be targeted through metabolic engineering to improve growth on the C1 substrate alone, bypassing the need for a second carbon source. As a first step toward this goal, in this work we conduct a thorough evaluation of co-metabolism in *E. limosum* across multiple conditions, comparing both C1-C1 and glucose-C1 cofeeding and significantly expanding upon previously tested substrate pairs. Specifically, we present the first characterization of CO and glucose co-feeding during formatotrophic growth in *E. limosum,* and the first evidence that CO co-feeding improves methylotrophic growth. *E. limosum* is a particularly attractive candidate for C1 bioconversion because of its substrate flexibility and recently developed advanced synthetic biology tools (Shin *et al*., 2019; Jeong *et al*., 2020; Flaiz *et al*., 2021; Sanford and Woolston, 2022a, 2024; Sanford *et al*., 2024).

One of the challenges of implementing and studying cofeeding is the potential for diauxie, or carbon-catabolite repression, in which the cell consumes the preferred substrate while ignoring the C1 substrate (Loubiere *et al*., 1992; Hill *et al*., 2025). Indeed, in the *E. limosum* study referenced above, during batch culture on glucose and methanol, co-consumption of methanol only began once the glucose concentration dropped below 6 mM (Loubiere *et al*., 1992). One solution is a fed-batch system, in which the preferred substrate is pumped continuously to the reactor at a rate below the biological uptake rate, so that it never accumulates in the media (Park *et al*., 2019). However, in a lab setting, screening of different fed-batch cofeeding conditions is not easily scaled. More broadly, low-cost, high-throughput anaerobic reactor options are not yet widely available, especially for fed-batch or gas feeding configurations needed for substrate cofeeding experiments. Commercial parallel bioreactor systems (e.g. Eppendorf DASGIP) and fluidics-enhanced microplate-based systems (e.g. Danaher BioLector) are highly expensive, and often incompatible with H_2_ in anaerobic gas mixtures. Several low-cost open-source alternative parallel reactor systems have been developed in recent years, such as the Chi.Bio (Steel *et al*., 2020) and eVOLVER (Wong *et al*., 2018; Heins *et al*., 2019). These highly customizable systems provide continous growth monitoring (via online OD) and flexible fluidics paradigms across multiple reaction vessels, but so far have not been adapted to support the growth of strict anaerobes outside an anaerobic chamber. Given the limited space available in most gloveboxes, a benchtop-compatible version would be highly advantageous. Continuous feeding of anaerobic gaseous feedstocks is even more inaccessible, as this typically requires pre-set compressed cylinder compositions and specially designed anaerobic reactors, posing space and cost obstacles for testing many conditions. Benchtop reactor systems with continuous feeding of custom anaerobic gas mixtures and liquid fed-batch configurations would dramatically improve throughput for cofeeding experiments. They would also broaden the accesibility of anaerobic microbiological research beyond specialist labs, benefitting multiple research areas, like the gut microbiome.

To overcome the limited reactor options and facilitate the screening of anaerobic cofeeding conditions described here, we adapted the eVOLVER system into AneVO. AneVO is a fully anaerobic benchtop reactor system with the same core functionality as eVOLVER (fluidic, temperature, and stir rate control as well as continuous optical density measurement), but which can maintain strict anaerobic conditions for 300+ hours on the benchtop with either liquid or continuous gas feeding. For glucose-C1 cofeeding, we established a robust system for anaerobic liquid feeding. For experiments with C1 gases, we paired AneVO with a flexible custom-built mass flow controller (MFC) based gas mixing system that enabled us to test different gas compositions simultaneously. This work validates AneVO as a low-cost parallel anaerobic bioreactor system that can be used by multiple research communities, as well as provides new insights into co-substrate metabolism in an emerging model acetogen.

## Results

### Fed-batch AneVO Maintains Strict Anaerobic Conditions on the Benchtop

To enable the high-throughput, convenient, anaerobic fed-batch experiments needed to study substrate cofeeding in *E. limosum,* we adapted eVOLVER into AneVO (**Figure 1a**). We first selected and tested materials for the culture vials. Of the options tested, two styles were successful: a 40-mL glass vial with a threaded cap sealed using Viton rubber septa, both of which are commercially available; and a custom-made (Chemglass) glass vial closed using the 20 mm butyl rubber septa and aluminum crimp seals used in traditional anaerobic microbiology with serum bottles. Both approaches maintained oxygen-free conditions for >144 hours, as indicated by the resazurin oxygen indicator remaining colorless (**Fig. 1b**). Part descriptions and sourcing are presented in **Table S1**, and further details regarding the selection and testing of the anaerobic media vial closures are presented in **Figure S1**. To validate AneVO against the current standard for acetogen lab culture, we compared batch growth of wild type *E. limosum* on methanol/CO_2_ in AneVO and serum bottles in parallel. The cell densities, growth rates, methanol uptake, and product concentration profiles were nearly identical between the AneVO vials and the serum bottles (**Fig. 1c**).

**FIGURE 1.**
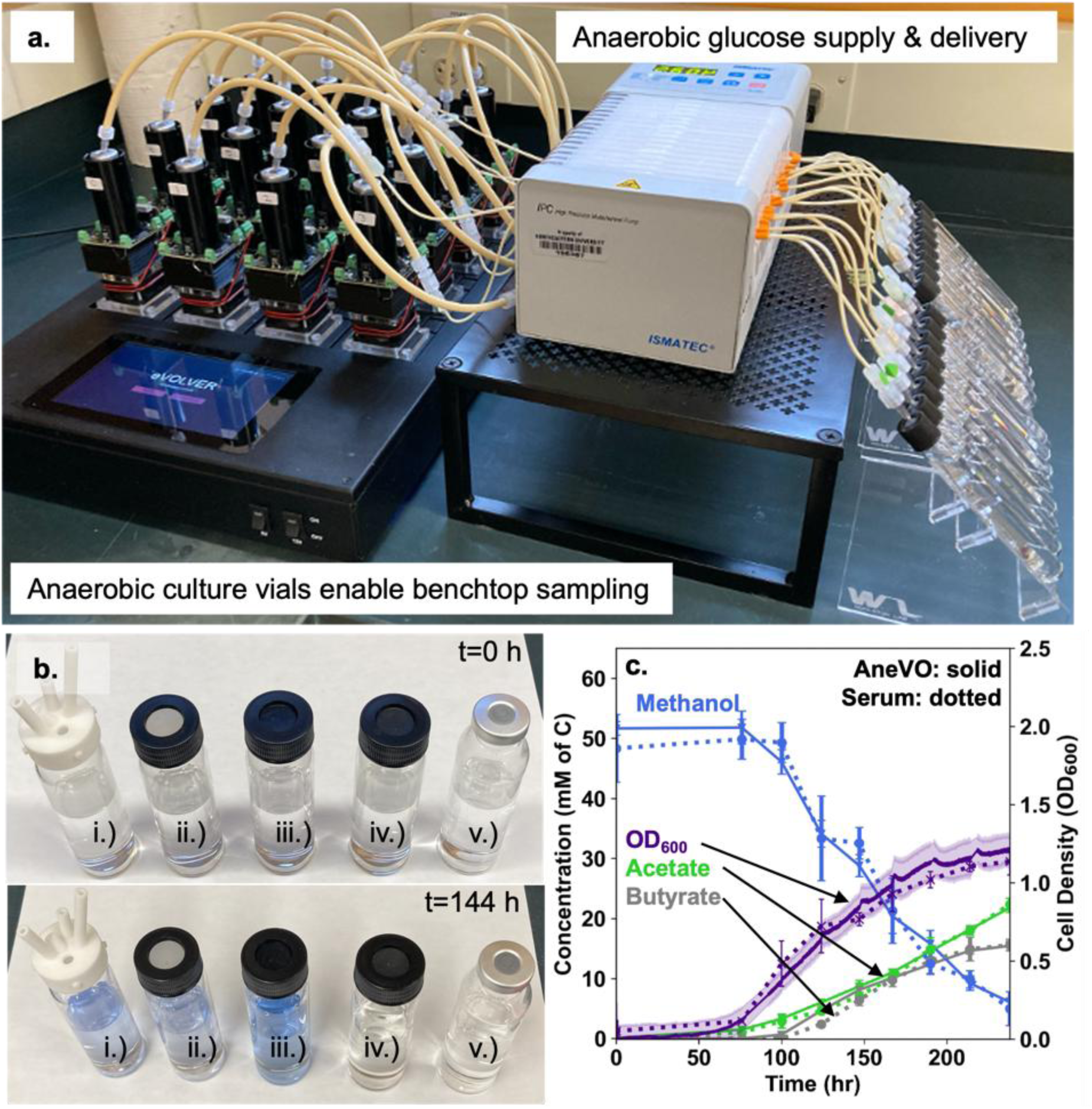
AneVO enables bench-top anaerobic growth experiments with continuous OD monitoring. a. Anaerobic glucose feeding was achieved using PharMed microbore tubing and needles through each septum. Separate pressurized Hungate tubes for each culture vial allows monitoring of glucose feed and maintenance of constant flow, all on the benchtop. b. Media vials were prepared in an anaerobic chamber using degassed vial components and media, before being brought to the benchtop. To test anaerobicity, the media contained colorimetric indicator resazurin and 3 mM of reducing agent cysteine. Materials of composition for each vial closure are as follows: i.) original eVOLVER silicon with nylon cap; ii.) silicon with threaded plastic cap; iii) hand-cut butyl rubber with threaded plastic cap; iv) Viton septa with threaded plastic cap; v) butyl rubber stopper with aluminum crimp top. c. AneVO was validated against serum bottles by comparing cell density, substrate uptake and product formation during growth of *E. limosum* on methanol and CO_2_.

To enable benchtop anaerobic fed-batch cofeeding experiments, we next developed a liquid feeding system for AneVO. To achieve this, we connected pressurized Hungate tubes to the culture vials using PharMed tubing with low oxygen permeability, needles and Luer fittings (**Fig. 1a** and **Table S1**), using a 16-channel peristaltic pump for precise dosing. In pilot growth experiments, both the culture vials and the glucose feed tubes remained anaerobic throughout the experiment, as confirmed by resazurin indicator and cell growth (data not shown).

### Glucose cofeeding improves growth and acetate productivity

We then used fed-batch AneVO to cofeed *E. limosum* with methanol and glucose (**Fig. 2**), as well as formate and glucose (**Fig. S2**). To avoid diauxie and ensure substrate co-metabolism, we inoculated AneVO vials containing the C1 substrate, and then began feeding 0.1 mM/h of glucose once the cell density began to increase. For both C1 substrates, growth benefited substantially from glucose cofeeding, with final cell density increasing 72% compared to methanol alone, and 254% compared to formate alone (**Fig. 2a, Fig. S2a, Table 1**). Throughout growth, the fed glucose was fully consumed and did not accumulate in the media. Combined with the continual depletion of the C1 substrates during the same timeframe (**Fig. 2b, Fig. S2b**), this shows that both substrates were metabolized simultaneously, and the improved growth was not simply from cells switching to consuming the preferred substrate.

**FIGURE 2.**
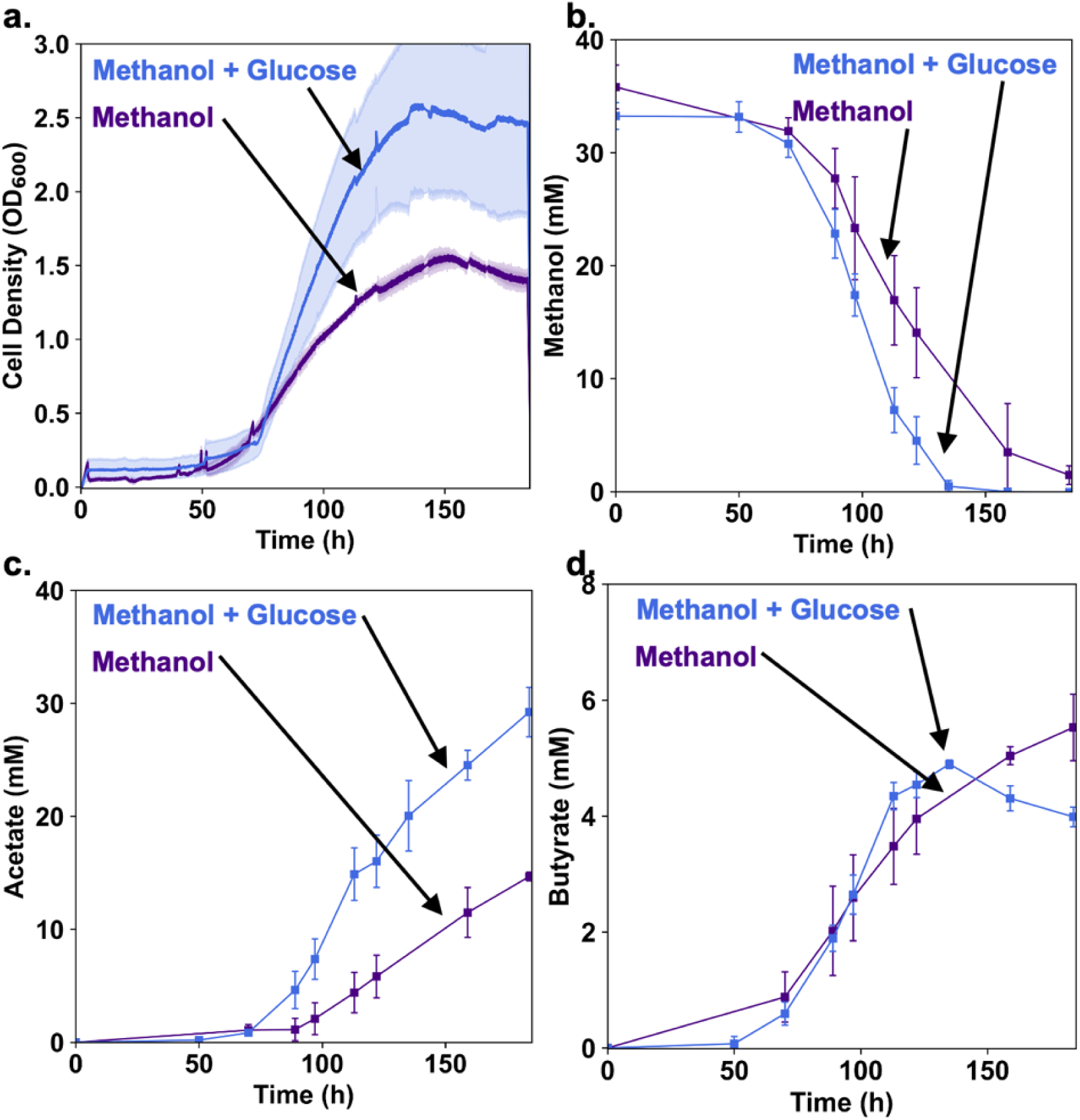
Growth and acetate productivity improved by methanol-glucose co-metabolism in fed-batch AneVO. Glucose was fed at 0.1 mM/h. a. Cell density measured as OD_600_ values in AneVO; b. Methanol concentration; c. Acetate concentration; d. Butyrate concentration. Plots display biological triplicates and error bars show standard deviation.

**Table 1.** Growth parameters for cofed *E. limosum* grown in AneVO. Mixotrophic and C1-C1 substrate pairs all lead to growth improvements, but vary in substrate uptake and acetate productivity outcomes. Final cell densities were calculated while primary C1 substrate remains. Media Conditions: G = glucose fed at 0.1 mM/h; CO = carbon monoxide; HCO3^-^ = sodium bicarbonate; HS = head space. Each culture vial included 0.05% yeast extract except for methanol v. methanol/glucose cultures. Formate v. formate/glucose cultures also included 10 mM sodium acetate.

| Primary C1 Compound | Cofeed Condition | CO<sub>2</sub> Supply | Maximal C1 Uptake<br>Q<sub>s</sub><br>mM/h | Maximal Specific C1 Uptake<br>q<sub>s</sub><br>mM/h/gDCW | Maximal Acetate Productivity<br>Q<sub>Ace</sub><br>mM/h | Maximal Specific Acetate Productivity<br>q<sub>Ace</sub><br>mM/h/gDCW | Final Cell Density<br>OD<sub>600</sub> |
| --- | --- | --- | --- | --- | --- | --- | --- |
| 50 mM Methanol | - | 50 mM HCO<sub>3</sub><sup>-</sup> | 0.62 ± 0.13 | 3.35 ± 0.63 | 0.20 ± 0.04 | 0.76 ± 0.14 | 1.32 ± 0.01 |
|  | G |  | 0.69 ± 0.08 | 3.25 ± 1.05 | 0.49 ± 0.11 | 1.34 ± 0.26 | 2.26 ± 0.13 |
|  | - | 14%-20% CO<sub>2</sub> constantly flushed through HS | 1.28 ± 0.23 | 10.52 ± 3.12 | 0.10 ± 0.01 | 1.73 ± 0.08 | 1.31 ± 0.18 |
|  | CO (15%) |  | 1.93 ± 0.15 | 5.25 ± 1.17 | 0.67 ± 0.16 | 3.39 ± 0.33 | 2.04 ± 0.09 |
|  | CO (30%) |  | 1.71 ± 0.14 | 5.19 ± 0.89 | 0.59 ± 0.10 | 2.11 ± 0.23 | 2.00 ± 0.04 |
| 100 mM Formate | - | 10 mM HCO<sub>3</sub><sup>-</sup><br>10% CO<sub>2</sub> HS | 1.79 ± 0.44 | 28.79 ± 3.36 | 0.59 ± 0.04 | 6.40 ± 0.70 | 0.54 ± 0.02 |
|  | G |  | 3.40 ± 0.39 | 21.43 ± 3.20 | 1.33 ± 0.83 | 5.94 ± 0.05 | 1.91 ± 0.14 |
|  | - | 14%-20% CO<sub>2</sub> constantly flushed through HS | 2.22 ± 0.29 | 26.74 ± 3.28 | 0.49 ± 0.06 | 5.85 ± 0.81 | 0.41 ± 0.01 |
|  | CO (15%) |  | 1.85 ± 0.11 | 11.99 ± 0.33 | 0.53 ± 0.04 | 2.90 ± 0.04 | 1.29 ± 0.17 |
|  | CO (30%) |  | 1.60 ± 0.27 | 5.28 ± 0.30 | 0.62 ± 0.07 | 1.96 ± 0.14 | 1.75 ± 0.05 |
| CO Only (30%) |  |  | - | - | 0.23 ± 0.01 | 1.33 ± 0.09 | 2.18 ± 0.08 |

As well as growth, acetate production from methanol benefited from cofeeding, with a 2-fold increase in final titer, and 2.4-fold increase in volumetric productivity over methanol alone (**Fig. 2c**, **Table 1**). Similarly, volumetric acetate productivity on formate improved 2.2-fold, though interestingly the final titer was no different than in the formate-only control (**Fig. S2c**). Butyrate was only produced from cultures fed with methanol, and titers were not impacted by methanol-glucose cofeeding. Interestingly, once the methanol was fully consumed, butyrate was partially consumed in the cofeeding condition (**Fig 2d**).

Overall methanol consumption was also faster during glucose feeding (**Fig. 2b**), although the maximal instantaneous uptake rate (both volumetric and specific) was not significantly altered (**Table 1**). On formate, glucose cofeeding resulted in a 90% increase in maximal volumetric uptake, but a 26% decrease in maximal specific uptake. Overall, glucose cofeeding improves growth, volumetric substrate uptake, and volumetric productivity on both formate and methanol, and product titer on methanol.

### Gas-Fed AneVO for C1-C1 Substrate Cofeeding Experiments

*E. limosum* is a particularly attrtactive candidate for substrate cofeeding because it can grow on a diverse range of C1 substrates, including CO. To be able to test gas-liquid substrate pairs like methanol/CO, we expanded AneVO to include continuous anaerobic gas feeding. To screen multiple gas compositions simultaneously, the gas-fed AneVO needed the capability to prepare strictly anaerobic gas mixtures at any desired ratio and serially feed the gas through the headspace of each culture vial. These design goals were met using a custom MFC-based gas mixing system (Alicat) equipped with an oxygen scrubber (**Figure 3**). Commercial pure gas cylinders feed a mixing tank at flow rates set by the master controller to generate the desired gas composition. From the mixing tank, mixed gas is routed through the oxygen scrubber (if needed), and is then delivered to individual rows of AneVO vials at a set rate by a dedicated mass flow controller. When necessary for specific experiments, additional gases can optionally be added into the mix downstream of the mixing tank for a specific set of vials with another mass flow controller. This has the benefit of allowing simultaneous testing of multiple different gas compositions, but with the downside of slightly diluting the other components, providing a compromise between flexibility and MFC cost. **Fig. 3** shows an example where for CO cofeeding where the tank is filled with a base CO_2_/N_2_ mixture, and CO is added to some vials at defined levels post-tank. All parts and sourcing are listed in **Table S1**.

**FIGURE 3.**
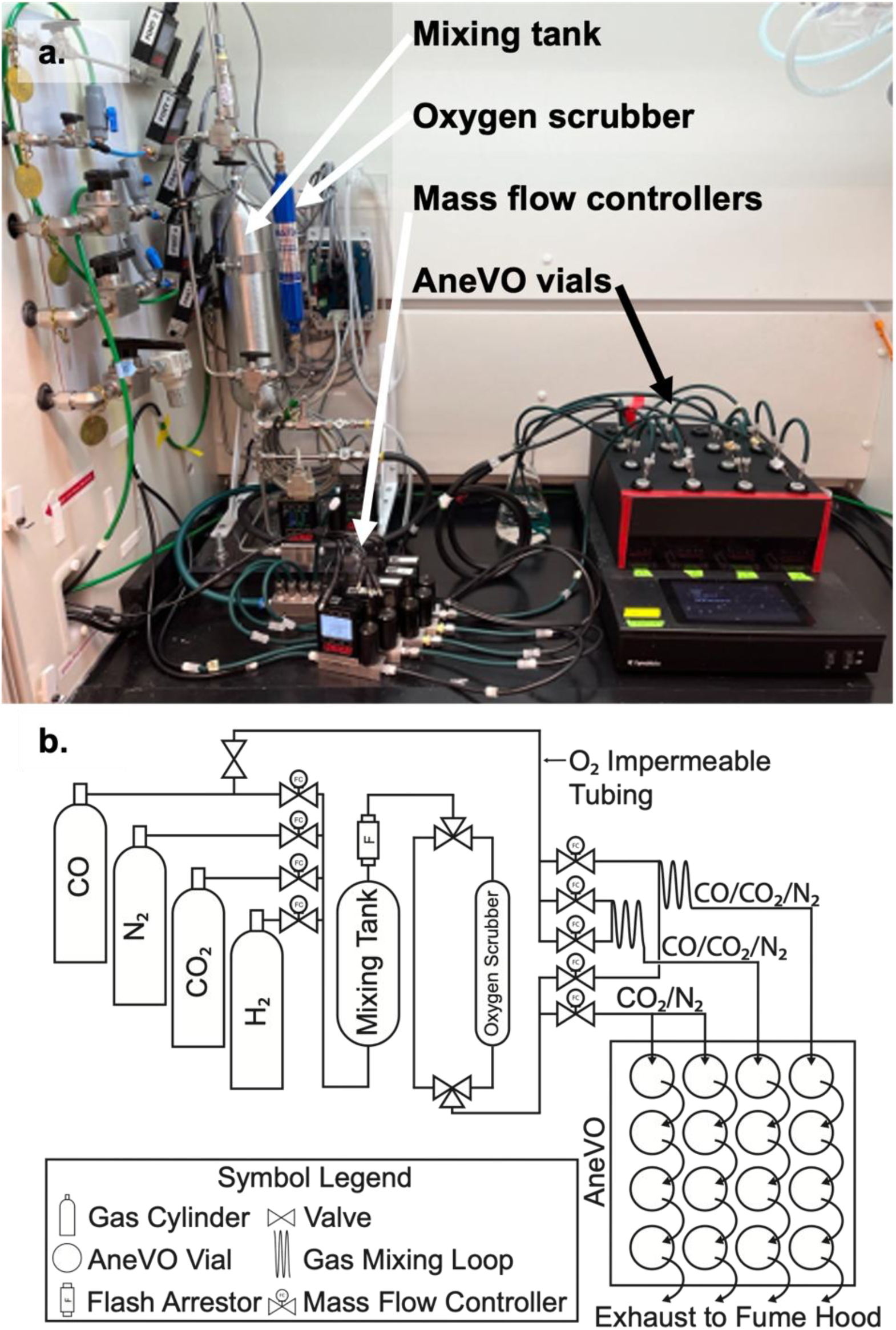
Continuous Gas Feeding in AneVO achieved with custom gas mixing system. a. AneVO includes continuous anaerobic gas feeding using a gas mixing system, with anaerobic septa, needles and anaerobic tubing. b. Piping and Instrument Diagram (P&ID) for continuous gas mixing system for AneVO.

### Methanol/CO Cofeeding Improves Growth Parameters Except Specific Uptake Rate

With the gas-fed AneVO set up, we grew *E. limosum* on methanol with no CO, 15% CO, and 30% CO (**Figure 4**), to evaluate the impact of CO cofeeding. In the cofed cultures, growth rate improved and final cell density increased by about 52% to 56% for either 15% or 30% CO respectively, compared to methanol alone (**Fig. 4a**). Similar growth between replicate cultures receiving gas in series (**Fig. 3b**) verified that the gas delivery rate was sufficient to maintain consistent gas headspace composition across vials, i.e. we infer that the gas flow rate was substantially higher than mass transfer rate to the liquid phase and/or the rate of cellular consumption. The maximum volumetric methanol uptake increased 50% and 34% for 15% CO and 30% CO, respectively and the maximal specific methanol uptake rate decreased by 50% for both CO feed rates (**Fig. 4b**, **Table 1**). The acetate productivity on methanol/CO was three-fold higher than the fed-batch methanol cultures with bicarbonate as the CO_2_ source, and five-fold higher than the gas-fed methanol cultures. The acetate titer also improved dramatically, by ∼5-fold (**Fig. 4c).** In the methanol/CO cultures, butyrate was partially consumed once the methanol was depleted, similarly to the methanol-glucose cofeeding condition (**Fig 4d**). Interestingly, there were differences in the methanol-only control in these experiments compared to the same control in the fed-batch experiments (**Table 1**). In the gas-fed experiments, fresh CO_2_ was constantly supplied through the headspace, while in the fed-batch experiments, sodium bicarbonate was continually supplied in the liquid feed. Both sets of cultures reached the same cell density of 1.3 OD_600_, but methanol uptake was two-fold higher in the gas-fed system. Acetate productivity was 50% lower in the gas-fed system relative to the fed-batch system.

**FIGURE 4.**
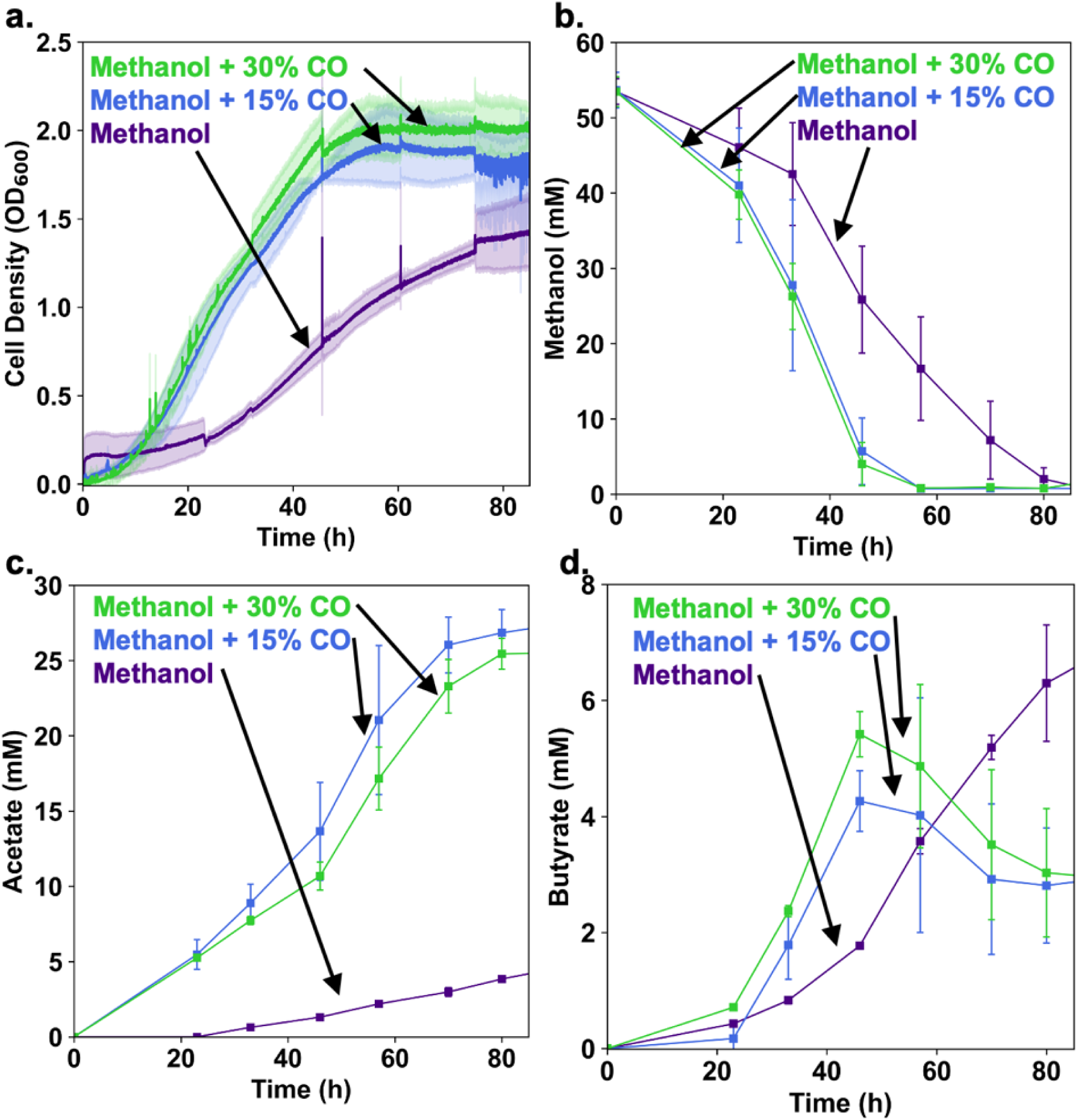
Methanol/CO cofeeding increases growth and product generation over methanol alone: 15% CO / 17% CO2 / 68% N2, 30% CO / 14% CO2 / 56% N2, and 20% CO2 / 80% N2, each at 10 SCCM. a. Cell density measured as OD600 values in AneVO; b. Methanol concentration; c. Acetate concentration. d. Butyrate concentration. Plots display biological triplicates and error bars show standard deviation.

### Formate/CO Cofeeding improves growth, but reduces specific acetate productivity

For comparison to methanol/CO cofeeding, we grew *E. limosum* on combinations of sodium formate with no CO, 15% CO, and 30% CO (**Fig. 5**). Growth on formate was significantly improved with CO cofeeding, with tripled and quadrupled cell density for 15% CO and 30% CO respectively (**Fig. 5a**). Growth on CO without any formate reached 4-fold higher than on formate alone (**Fig. 5a**).

**FIGURE 5.**
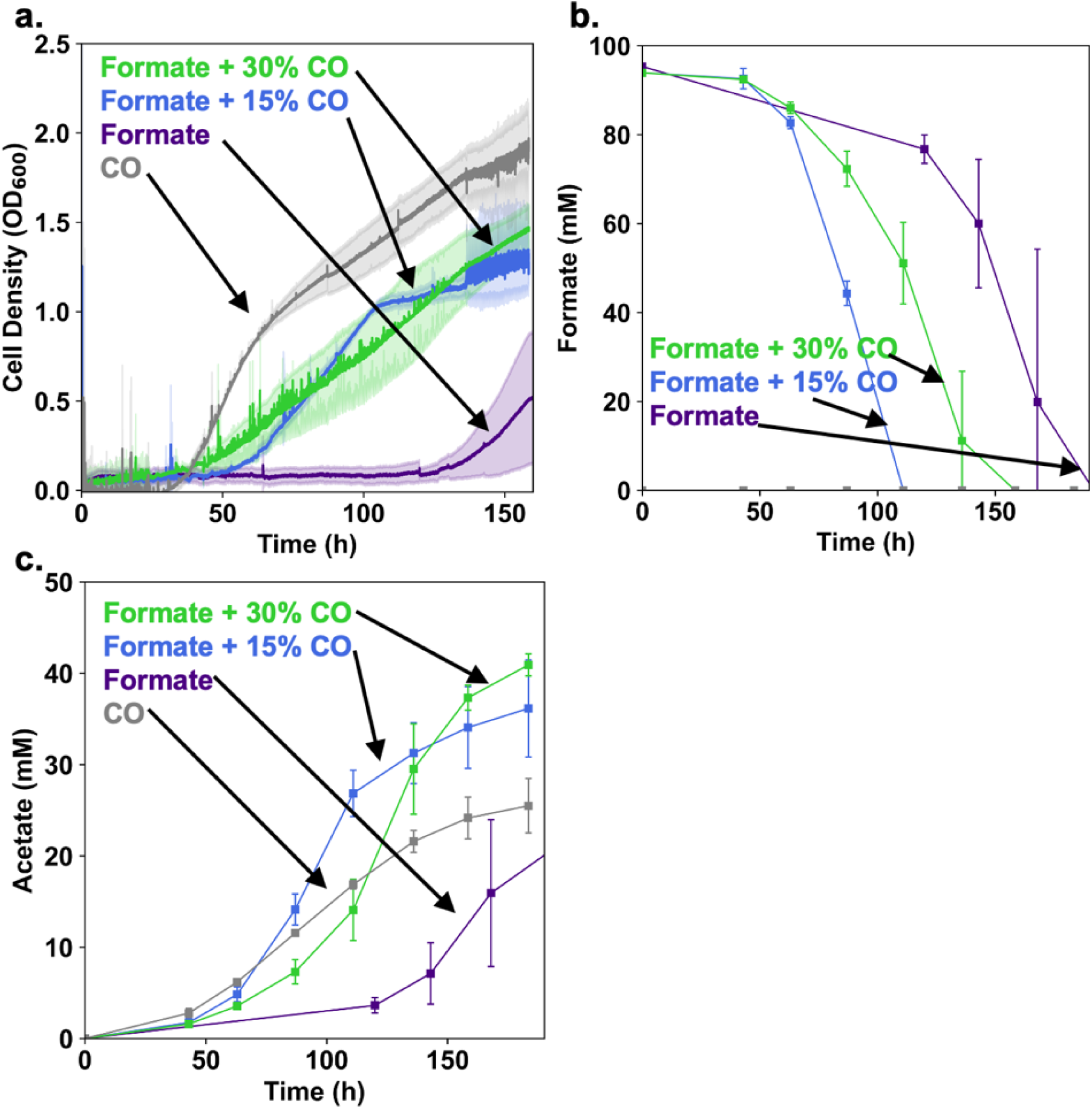
Formate/CO cofeeding improves growth and volumetric acetate productivity: 15% CO / 17% CO2 / 68% N2, 30% CO / 14% CO2 / 56% N2, and 20% CO2 / 80% N2, each at 10 SCCM. a. Cell density measured as OD600 values in AneVO; b. Methanol concentration; c. Acetate concentration. If butyrate was produced, it was below detection levels. Plots display biological triplicates and error bars show standard deviation.

Interestingly, the volumetric formate uptake rate remained unchanged when cofed CO, considering variation between replicates (**Fig. 5b**). The specific formate uptake decreased by 55% for the 15% CO condition, and 80% for the 30% CO condition. The decrease in specific uptake for the 15% CO condition was similar to that observed for methanol/CO cofeeding. Additionally, while the acetate titer improved substantially (**Fig. 5c**), the volumetric productivity increased only marginally, and the specific acetate productivity decreased by 50% and 66% for the 15% CO and 30% CO conditions, respectively.

## Discussion

Our focus in this work was to establish a convenient parallel anaerobic reactor system for fed-batch and gas fermentation experiments, then to use this platform to expand our knowledge of substrate cofeeding in *E. limosum,* with the aim of i) identifying promising substrate combinations for producing value-added products in engineered strains at higher rates and titers, and ii) providing a basis for future mechanistic investigation into how co-feeding improves fitness. AneVO enables benchtop anaerobic cultivation under batch and fed-batch conditions, using entirely low-cost commercially available parts (**Table S1**). Addition of a home-built MFC system for gas mixing enables continuous feeding of anaerobic gas mixes at a fraction of the cost of custom gas mixes or turn-key gas mixing systems. This system allowed us to rapidly test co-feeding of both liquid substrates (glucose) and gaseous ones (CO). As acetogens are some of the most oxygen-sensitive microbes known, successful cultivation of *E. limosum* suggests the system could be used for a wide range of strict and facultative anaerobes.

Our co-feeding results with glucose/methanol mixtures confirm previous observations from chemostat cultures (Loubiere *et al*., 1992) and extend them to fed-batch operation. Glucose co-feeding during formatotrophic growth has not been previously evaluated in *E. limosum,* and resulted in an even greater increase in biomass concentration than during methylotrophic growth (254% vs 72%). Most excitingly, CO supplementation dramatically enhanced both formatotrophic and methylotrophic growth, with a > 5-fold improvement in acetate titer on methanol (**Fig. 4c**). While previous work showed that methanol improved growth on CO in *E. limosum* KIST612 compared to controls without methanol, whether CO boosted growth on methanol was not determined (Kim *et al*., 2021). Here we found a substantial improvement in volumetric methanol uptake rate with CO supplementation. CO is formed alongside methanol and/or formate in electrocatalytic systems for CO_2_ conversion (Schwarz *et al*., 2024), thus *E. limosum’s* improved growth from simultaneous consumption of CO and methanol may provide a particularly attractive opportunity to both better utilize these impure feedstocks, and overcome the energy limitations of acetogenesis on a single C1 substrate.

From a mechanistic standpoint, there are several possible hypotheses to explain the fitness benefit from co-feeding. In acetogens, pyruvate:ferredoxin oxidoreductase (PFOR) connects the WLP to gluconeogenesis for generating biomass precursors, fixing CO_2_ and consuming reduced ferredoxin (Fd_red_) during autotrophic growth (Furdui and Ragsdale, 2000; Song *et al*., 2018). PFOR is one of the most abundant proteins in *E. limosum* (Song *et al*., 2018), and the reaction it catalyzes is thermodynamically challenging in the reductive direction, suggesting it may be rate-limiting. Since methanol and formate are metabolized via the WLP, and glucose via glycolysis, one possibility is that glucose cofeeding reduces or eliminates the need for reductive flux through PFOR, thus improving growth (Le Bloas *et al*., 1993). Confirming this would require ^13^C-metabolic flux analysis (MFA), the gold standard for quantifying carbon flux direction and magnitude (Long and Antoniewicz, 2019). ^13^C-MFA is often challenging in C1-consuming organisms grown on a single substrate because all carbon gets uniformly labeled, but cofeeding overcomes this challenge (Hoyt and Woolston, 2022). Additionally, the AneVO system developed here is ideal for isotopic labeling studies because of the small volume (15-40 mL) of culture media required.

Alternatively, the improved growth rate and cell density may be due to increased ATP availability. Growth on C1 compounds using the WLP is efficient, but provides <1 ATP per acetate (Schuchmann and Müller, 2014). By contrast, conversion of hexoses to acetate through glycolysis provides 4 ATP. Indeed, the acetate titer during methanol cofeeding increased by 2.2-fold compared to methanol. If this extra flux came from glucose, this would represent a significant increase in ATP availability. Interestingly, although glucose cofeeding substantially improved growth on formate, the acetate titer did not significantly increase (**Fig. S2**), potentially suggesting an alternative mechanism in this case. This result is also in contrast to similar experiments in *A. woodii* where fructose supplementation resulted in an acetate titer of 16.2 g/L, compared to 3.17g/L on formate alone (Neuendorf *et al*., 2021). This could imply that the mechanisms underlying the improved fitness are unique to individual acetogens.

The benefits of cofeeding could also be due to improved production of reduced ferredoxin (Fd_red_). Uptake of CO is catalyzed by the reversible carbon monoxide dehydrogenase (CODH), producing Fd_red_ and CO_2_ (Dietrich *et al*., 2021; Kremp and Müller, 2021), as shown in **Figure S3**. Likewise, oxidative flux through PFOR from glucose produces Fd_red_. Ferredoxin is the electron carrier with the greatest reducing potential in acetogens, and is crucial for acetogenic growth on any C1 substrate, where it is used in the reductive direction by PFOR, as well as for energy conservation by the Rnf or Ech complexes (Schuchmann and Müller, 2014). Additionally, reduced ferredoxin is thought to be required for methanol oxidation by methylene tetrahydrofolate reductase (MTHFR) in *E. limosum* (Dietrich *et al*., 2021). CO cofeeding was especially beneficial for methanol conversion, consistent with the possibility that the high demand for Fd_red_ on this substrate is the limits growth on methanol alone. Supply of CO would also feed the carbonyl branch of the WLP, obviating the need for reduced ferredoxin to assimilate CO_2_ for acetyl-CoA generation.

Alternatively, the benefit of co-feeding could stem from routing carbon flow away from flux-controlling reactions. During growth on methanol and formate, a portion of the substrate is oxidized through the methyl branch of the WLP to provide reducing equivalents, while the rest is incorporated into the methyl group of acetyl-CoA (Moon *et al*., 2021). It is possible that one of these reactions is kinetically or thermodynamically limited, and that providing additional reducing equivalents through CO or glucose requires less oxidative methyl branch flux, overcoming a rate-controlling step and resulting in faster growth. Again, determination of how these fluxes vary between mixotrophy and single-substrate growth with ^13^C-MFA would help identify the underlying mechanism.

A universal observation in our work was that the specific uptake of the primary C1 substrate decreased during cofeeding with either glucose or CO. While Neuendorf and coworkers also observed specific C1 uptake decreases for formate/CO and formate/fructose substrate pairs in *A. woodii* (Neuendorf *et al*., 2021), in other cases cofeeding has resulted in the opposite effect, with a synergistic effect on growth and C1 substrate uptake. The Lindley group observed a 33% increase in specific methanol uptake rate in a different *E. limosum* strain for a short period in the middle of batch growth, and a maximal dilution rate in chemostat experiments that was higher during co-feeding than on either glucose or methanol alone (Loubiere *et al*., 1992). In *M. thermoacetica*, glucose cofeeding significantly improved specific CO_2_ uptake (Park *et al*., 2019). Finally, Wood and coworkers saw 2.5 fold increase in specific methanol uptake in *E. limosum* when formate was co-fed at a 1:3 ratio in chemostats (Wood *et al*., 2023). Why the outcome is different across acetogens remains unclear, as does whether the decrease is the outcome of a deliberate regulatory system or a metabolic bottleneck. Elucidating the underlying mechanisms would be a particularly compelling use of the expanding genetic and systems biology tools for *E. limosum.* If this limitation is indeed regulatory, then cells could in principle be engineered to maintain the maximal specific C1 uptake rate while benefitting from co-metabolism, leading to even further improved performance metrics.

## Conclusions

*E. limosum* is rapidly emerging as a model acetogen for production of biofuels and biochemicals from sustainable C1 feedstocks, thanks to its substrate flexibility and genetic tractability. The cofeeding experiments of this study significantly expand our knowledge of C1 metabolism in this species. They also suggest several strategies for enhancing C1 bioconversion metrics in *E. limosum,* in particular the simultaneous use of methanol and CO to enhance substrate uptake, growth, and product titer. With these baseline phenotypic data established, future studies should focus on elucidating the underlying mechanisms through a combination of ^13^C isotopic tracer analysis and genetics approaches, and understanding why the specific C1 uptake rate decreases. The expanding genetic toolkit for *E. limosum* may offer a means to deregulate substrate uptake, or use the energy available from cofeeding to make higher value products. More broadly, the AneVO platform developed and validated here provides a low-cost system for moderate throughput benchtop anaerobic culturing experiments in batch, liquid fed-batch, and continuous gas-fed configurations. A full parts list is available in the supplement, facilitating its deployment for a wide range of anaerobic microbiology experiments. As research interest in anaerobes continues to grow, motivated by applications in human health and sustainability, we believe enhanced access to convenient reactor systems such as AneVO for handling these oxygen-sensitive organisms in different fluidics modes will enable broader participation of scientists in this exciting work.

## Experimental Procedures

### Bacterial strains, media and growth conditions

Growth experiments were conducted with wild type *E. limosum* ATCC 8486 from the German Collection of Microorganisms and Cell Cultures (DSMZ). Cells were stored in cryogenic stocks at -80°C, and then precultured in the medium matching the experimental conditions. Low salt modified Gottschalk minimal medium, a fully-defined low salt medium (“basal medium”), was prepared in a Coy anaerobic chamber and supplemented with varying carbon sources for each growth experiment (Kountz *et al*., 2020). The media salts in the basal medium were comprised of: 1 g/L NH_4_Cl; 0.1 g/L MgSO_4_•7H_2_O; 50 mg/L CaCl_2_•2H_2_O, 2.25 g/L NaCl. The Eubacterium Vitamins were included per liter as follows: 0.1 mg of biotin, 0.2 mg of folic acid, 0.3 mg of pyridoxine-HCl, 0.2 mg of thiamine-HCl, 0.1 mg of riboflavin, 0.2 mg of nicotinic acid, 0.2 mg of D-pantothenate, 0.1 mg of vitamin B12, 0.1 mg of p-aminobenzoic acid, and 0.1 mg of lipoic acid. Pfennig and Lippert’s trace elements solution (Pfennig and Lippert, 1966) was included at 3 ml/L. Selenite and tungstate solution was used at 0.5 mg/L of NaOH, 3 μg/L of Na_2_SeO_3_•5 H_2_O, and 4 μg/L of Na_2_WO_4_ • 2 H_2_O. A concentration of 3 mM HCl-cysteine was used to reduce the media, with 1 mL/L of 0.1% Resazurin indicator. 50 mM potassium phosphate buffer was included, and the media was adjusted to pH 7.5.

Media conditions tested included either 50 mM methanol or 100 mM sodium formate as the primary carbon source. Methanol culture medium also included 50 mM sodium bicarbonate and was fully defined for the glucose cofeeding experiment. Formate culture medium included 10 mM sodium bicarbonate, 10 mM sodium acetate, and 0.05% yeast extract for the glucose cofeeding experiment. All gas-fed culture media included 0.05% yeast extract.

### Batch Cultivation in AneVO

40-mL custom glass vials were prepared with 25-mL media volumes in an anaerobic chamber with an atmosphere of 10% CO_2_, 5% H_2_, and 85% N_2_, except for methanol v. methanol/glucose which was 5% H_2_, and 95% N_2_. Vials were sealed in the anaerobic chamber and brought to the benchtop for use in the eVOLVER vial platform. Per the procedures originally established for eVOLVER, a magnetic stir bar was put in each vial prior to sealing, to keep cells suspended for consistent OD monitoring. Both styles of glass vial were produced by Chem Glass (Vineland, New Jersey).

### Fed-Batch Cultivation in AneVO

The feed system was designed for low flow rates so that, in combination with removed sample volumes, the culture volume remained approximately constant over the duration of the experiment. Hungate tubes for glucose cofeeding were filled with basal medium containing 33.6 mM of glucose, and the volumetric feeding rate was 1.29 μL per minute, for a feed rate of 2.6 μmol per hour. PharMed tubing, Luer fittings, and needles were connected, autoclaved, and then purged with sterile anaerobic media prior to connection with the Hungates and culture vials. For fed-batch growth conditions, cultures were inoculated at 4% v/v into the C1 substrate, and once the OD reached approximately 0.1, flow of glucose was initiated with an Ismatec IPC High Precision Multichannel Pump and maintained continuously until the OD was no longer increasing.

Benchtop sampling followed classic anaerobic microbiology techniques, by purging the syringe and needle with anerobic gas, and replacing the volume of gas removed from the culture vial, prior to collecting a sample or inoculation. 600 μL culture media samples were collected for each time point, centrifuged at 4 °C, 20,000 RCF for 10 minutes, and then the supernatant was frozen in tubes until analysis.

### Gas-Fed Cultivation in AneVO

Using the same media and vials as the fed-batch cultivation, the vials were prepared with media and closed, but for gas feeding the septa were pierced with two 22- gauge 1.5 inch needles, one for the inlet gas feed and another for the outlet. The needles were kept above the media level ensuring the OD readings were not disturbed by bubbling and to minimize evaporation. The vials were fed in series, with the efflux piped to the next sequential culture vial with the same gas composition, for up to a maximum of four vials. The gas flow rate (10 SCCM) was calculated to deliver CO at a rate 10-fold higher than the consumption rate for autotrophically growing cells at a maximal OD_600_ of 2.0 (see **Supplementary Note S1**), such that even neglecting mass transfer resistance, there would be no more than a 10% change in gas composition between sequential vials. Given the poor solubility of CO and low mass transfer coefficient in the absence of sparging, the composition change is likely much less than this. The gas feeding levels used here included: 20% CO_2_ / 80% N_2_, 15% CO / 17% CO_2_ / 68% N_2_, and 30% CO / 14% CO_2_ / 56% N_2_, each at 10 SCCM.

### Analytical methods

Concentrations of glucose, acetate, and butyrate in culture medium were quantified by High Pressure Liquid Chromatography (Agilent 1260 Infinity II), equipped with a BioRad Aminex HPX-87H ion exchange column (300mm × 7.8 mm) at 60 °C, and a refractive index detector at 50 °C. 10 μL injections were analyzed with an isocratic method using 14 mM sulfuric acid at a flowrate of 0.6 mL/min, and concentrations determined by comparing peak areas to a standard curve (Hayes *et al*., 2023).

Methanol concentration was quantified using a modification of a colorimetric enzymatic assay in 350 μL, 96-well plate (Anthon and Barrett, 2004). First the samples were diluted into the linear assay range with water, while alcohol oxidase (from *Pichia pastoris,* E.C.1.1.3.13 from MP Biomedicals) was prepared at a concentration of approximately 5 units per mL of 7.5 pH, 100 mM sodium phosphate buffer. To perform the well-plate assay, 20 μL of diluted sample was added to 80 μL of water, treated with 100 μL of the alcohol oxidase in buffer, then incubated for 10 minutes at 30 °C which produced formaldehyde and hydrogen peroxide. The Purpald (4-Amino-3-hydrazino-5-mercapto-1,2,4-triazole from Alfa Aesar) was freshly prepared at 5 mg/mL in 0.5N sodium hydroxide solution, then diluted 50% by volume with water. 150 μL of the Purpald mixture was added to each assay well, and the plate was incubated for 30 additional minutes at 30 °C. The resulting purple tetrazine, proportional to the original methanol concentration, was detected by measuring absorbance at 550 nm in a Molecular Devices SpectraMax i3 plate reader. A standard curve of methanol in matching basal medium and dilution was prepared in each plate, ranging from 0 mM to 50 mM.

### Calculation of growth parameters

The substrate uptake rates and product generation rates were calculated using the optical density and supernatant concentration data. The OD:Dry Cell Weight (DCW) ratio was assumed to be 0.3 gDCW/L per OD-L(Loubiere *et al*., 1992). C1 uptake was calculated from the change in concentrations at each time point, with specific uptake calculated using the average cell density for the time range. The maximal C1 uptake value was calculated from the highest uptake rate for each replicate culture vial regardless of point in time, and averaged together for the same growth condition.

Acetate productivity and specific acetate productivity were also calculated for each time point, using the average cell density for the time range, with the highest value for each vial averaged together for each condition. For formate v. formate/glucose, which included 10 mM sodium acetate in the initial media, the net produced acetate was used.

## Acknowledgements

This work was supported by Startup Funds from Northeastern University, the DOE ARPA-E ECOSynBio program (Award #DE-AR0001511), the National Science Foundation (Award #2345872), the National Institutes of Health (NIH) (grant nos. R01AI171100, R01EB027793), and the Schmidt Sciences Polymath award (no. G-22 63292). KOH was additionally supported by the Ford Foundation Predoctoral Fellowship. We thank Brandon Wong at FynchBio and Boston University for the eVOLVER collaboration, and Seung Hwan Lee at Massachusetts Institute of Technology for helpful discussions about energy cofactor balances. We thank the Snell 3D Printing Studio Team for the assistance with generating the tube holders, and Frank Lacivito and the team at Chemglass for creating the custom glass vials.

## Supporting Information

All data supporting the conclusions of this work are contained in the main text and supplementary material. A detailed step-by-step protocol for methanol detection via the Purpald Assay is available at the Woolston lab website: www.woolstonlab.org.

## Notes

### Competing Interest Statement

A.S.K. is a scientific adviser for and holds equity in Senti Biosciences and Chroma Medicine, and is a cofounder of K2 Therapeutics and Fynch Biosciences, a manufacturer of eVOLVER hardware. All other authors declare no conflicts of interest.

