## Supplemental Information for "Exploring C1 substrate cofeeding in *Eubacterium limosum* with AneVO, a low-cost anaerobic parallel bioreactor platform"

**Table S1. AneVO Parts and Sourcing Information**

| Category | Component | Part | Supplier and Part Number (PN) |
|---|---|---|---|
| Media Vials | 40-mL glass vial with screw top closure, suitable for liquid media | 40 mL glass vial with screw top, 28mm x 95mm | Chemglass, PN CG-4902-08 |
| | | Polypropylene Screw Cap with Hole 24-mm | Supelco PN 27057 |
| | | Viton Septa, 60 Mils Thick, 0.866" Diameter | Supelco PN 27355 |
| | 40-mL glass vial with crimp top closure, suitable for liquid or gas fed media | Custom glass vial with crimp top closure | Chemglass<br>PN NOR-2104-061CV |
| | | 20 mm butyl rubber septa and aluminum crimp tops made for Wheaton bottles | Stopper PN 11-102-0929 (Fisher);<br>Crimp top PN 11-100-5224 (Fisher) |
| Media Delivery | Peristaltic pump and compatible microbore tubing | Either 4-roller or 16-roller, low flow pump, and Ismaprene microbore tubing 400 mm length, with 0.25 mm ID | Ismatec IPC High Precision Multichannel Pump (16 roller);<br>PharMed 2 stop tubing PN 95723-12 |
| | | | Ismatec Reglo ICC Pump (4 roller);<br>Ismatec PharMed BPT 3 stop tubing PN 95714-12 |
| | Microbore fittings | 0.010 " ID to connect microbore tubing to male luer connector, additional Luer fittings as needed. | Component Supply<br>Part Number MBF-1-MLL |
| | Hungate tubes, septa, screw caps | Stopper is Butyl Rubber, 5.4mm;<br>Screw cap is 9 mm open top;<br>Hungate tube is 16 x 125mm | Stopper PN 50-121-5190 (Fisher);<br>Screw cap PN 50-121-5189 (Fisher); Full set PN 50-121-5187 (Fisher) |
| | Sterile needles with Luer connection | 22G x 1.5 " for vial feed and sampling;<br>optionally 22G x 3 " for sampling;<br>22G x 4 " for glucose delivery tubes | Varied |

**Table S1 (Continued). AneVO Parts and Sourcing Information**

| Category | Component | Part | Supplier and Part Number (PN) |
|---|---|---|---|
| Gas Feeding | Gas feeding tubes | 0.25" ID 0.5" OD neoprene rubber tubing | McMaster-Carr PN 5034K21 |
| | | 0.25" barb luer lock fittings (nylon) | varied |
| | Sterile needles with Luer connection | 22G x 1.5 " for gas inlfux and efflux | Varied |
| | Mass Flow Controller (MFC) | Higher flux MFC | Alicat - MCW Series Controller 10 - 100 SCCM |
| | | Lower flux MFC | Alicat - MCW Series Controller 0.5 - 5 SCCM |
| Gas Mixing | Mass Flow Controller mixing setup | Input feed MFC, controller, and controller board | Alicat designed and supplied |
| | 1/4 stainless steel tubes | For all high pressure gas lines and for non-flexible connections | varies |
| | Swagelok fittings | For all high pressure stainless steel tubing connections | varies |
| | Mixing tank | Metal tank for mixing various gas compositions | Swagelok 304L-HDF4-2250 |
| | Oxygen scrubber | Scrubber to achieve inline removal of oxygen from gas mixes | Restek 20600, Fisher Scientific PN 06-710-843 |
| | Flash arrestor | To prevent any sparks from returning to flammable gas supply tanks | Varies |
| | One way check valve | To prevent pressure build up from returning to gas cylinders | Varies |

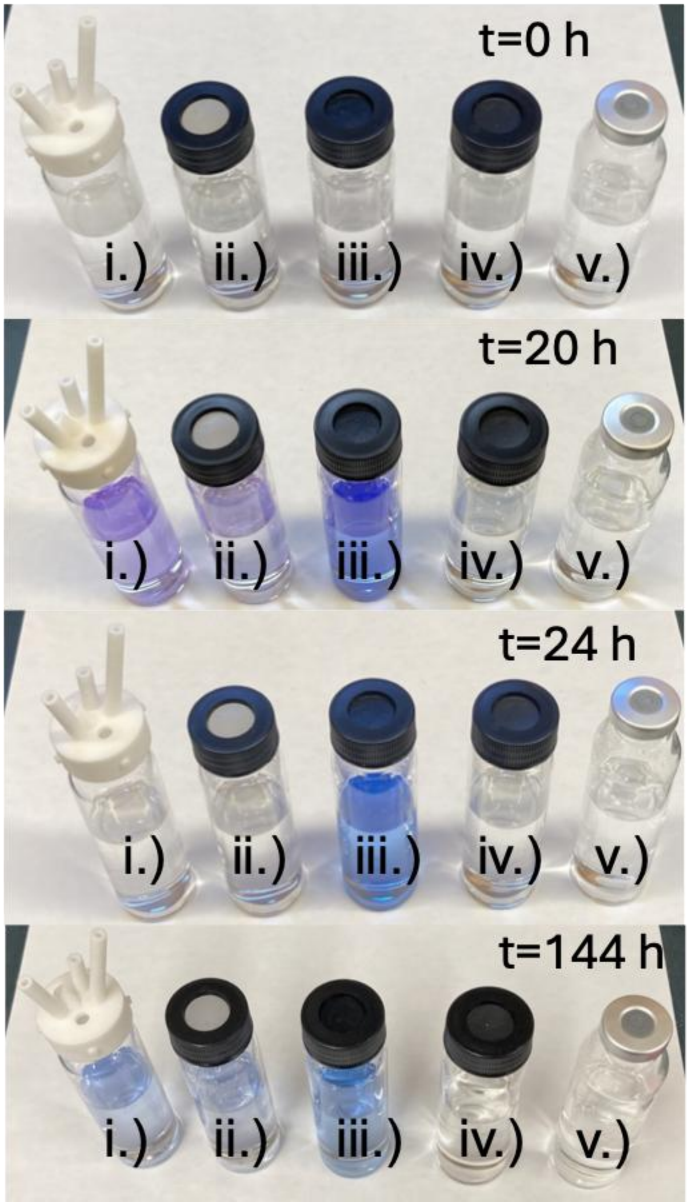


**Figure S1. Anaerobic vial closure testing and selection.** Media vials were prepared in an anaerobic chamber using degassed vial components and media, before being brought to the benchtop. To test anaerobicity, the media contained colorimetric indicator Resazurin and 3 mM of reducing agent cysteine. Materials of composition for each vial closure are as follows: i.) original eVOLVER silicon with nylon cap; ii.) silicon with threaded plastic cap; iii) hand-cut butyl rubber with threaded plastic cap; iv) Viton septa with threaded plastic cap; v) butyl rubber stopper with aluminum crimp top.

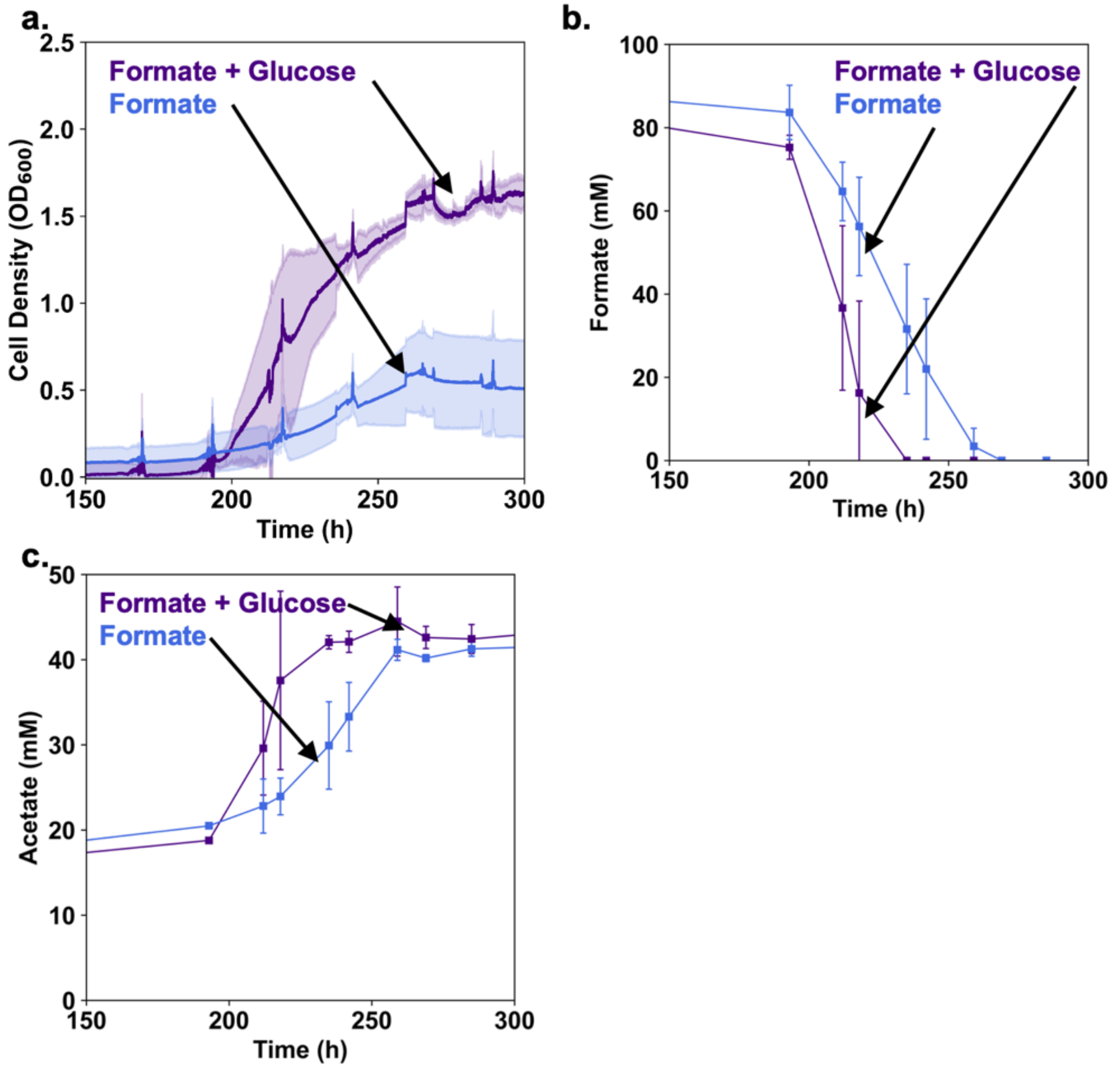


**Figure S2. Growth of *E. limosum* on formate as primary substrate with and without glucose cofeeding in fed-batch AneVO.** Glucose feeding was 0.01 mM/h. Each culture vial included 10 mM sodium acetate, 10 mM sodium bicarbonate, and 0.05 % yeast extract. a.) Cell density measured as OD600 values in AneVO. b.) Methanol concentration measured via enzymatic assay. c.) Acetate values measure with HPLC. If butyrate was produced, it was below detection levels. Plots display biological triplicates for growth on formate alone, and duplicates for formate cofed with glucose. Error bars show standard deviation.

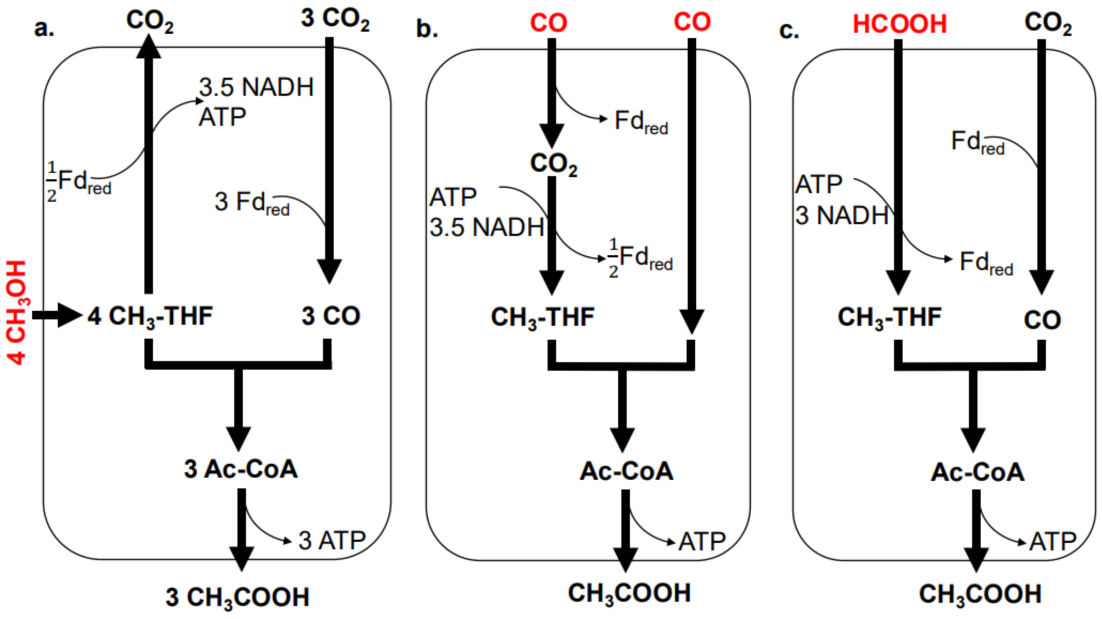


**Figure S3. Wood-Ljungdahl Pathway (or Reductive Acetyl-CoA Pathway) for *E. limosum* showing energy cofactor differences and changes in flux direction for each C1 substrate.** Chemiosmotic energy generation in *E. limosum* includes Rnf and ATP Synthase and allow exchange of energy cofactors (not shown). a.) Growth on methanol and $CO_2$. b.) Growth on carbon monoxide. C.) Growth on formate.

**Note S1. Gas Flow Calculations**

First calculate the maximal CO uptake rate as follows:

Maximal specific CO uptake rate (mmol/(g-h)) from (Hermann *et al.*, 2020)

$q_{CO} = 18.4$

Biomass Concentration (g/L), based on OD 2.0, and 0.25 g cell / OD-L)

$X = 0.5$

Reactor volume (L)

$V = 0.04$

Maximal CO uptake rate (mmol/h) is calculated:

$Q_{CO} = q_{CO}*X*V$

The Maximal CO uptake rate (per vial) is 0.37 mmol/h.

Then calculate CO delivery rate as follows:

Gas flow rate (L/h, with 10 mL/min), typically used for AneVO gas feeding system:

$F = 10 * 60 / 1000$

Mol fraction of CO in gas mix

$y_{CO} = 0.15$

Pressure (atm)

$P = 1$

Temperature (K)

$T = 37+273.15$

Gas Constant (L-atm/mol-K)

$R = 0.082$

Gas Molar Volume (L/mol)

$v = R*T/P$

CO delivery rate (mmol/h) is calculated:

$CO_{delivery\ rate} = F*y_{CO}/v*1000$

The CO delivery rate is 3.54 mmol/h for 15% CO gas fed at 10 mL/minute, 10-fold higher than the maximal CO uptake rate.
